# Energy metabolites mediated cross-protection to heat, drought and starvation induced plastic responses in tropical *D.ananassae* of wet-dry seasons

**DOI:** 10.1101/158634

**Authors:** Chanderkala Lambhod, Ankita Pathak, Ashok K Munjal, Ravi Parkash

## Abstract

Cross-tolerance effects for cold and drought stressors are well known for temperate and polar ectothermic organisms. However, less attention has been paid to plastic changes induced by wet-dry conditions for the tropical insect taxa. *Drosophila ananassae* is abundant in wet habitats but its desiccation sensitivity is likely to make it vulnerable under expected drought conditions due to climate change. We tested plastic effects of heat hardening, acclimation to drought or starvation; and changes in trehalose, proline and body lipids in *D. ananassae* flies reared under wet or dry season specific conditions. Wet season flies showed significant increase in heat knockdown, starvation resistance and body lipids after heat hardening. However, accumulation of proline was observed only after desiccation acclimation of dry season flies while wet season flies elicited no proline but trehalose only. Thus, drought induced proline can be a marker metabolite for dry season flies. Further, partial utilization of proline and trehalose under heat hardening reflects their possible thermoprotective effects. Heat hardening elicited cross-protection to starvation stress. Stressor specific accumulation or utilization as well as rates of metabolic change for each energy metabolite were significantly higher in wet season flies than dry season flies. Energy budget changes due to inter-related stressors (heat vs desiccation or starvation) resulted in possible maintenance of energetic homeostasis in wet or dry season flies. Thus, low or high humidity induced plastic changes in energy metabolites can provide cross-protection to seasonally varying climatic stressors.

**Summary statement:** In the tropical *Drosophila ananassae*, low or high humidity induced plastic changes in energy metabolites provide cross-protection to seasonally varying climatic stressors

## INTRODUCTION

In temperate regions, ectothermic organisms encounter a greater range of colder environments (freezing to mild warm), as well as changes in other abiotic factors which affect their acclimatization to improve survival under harsh climatic conditions (Angilletta, 2009; Sinclair, 2015). Ectothermic organisms from temperate or polar regions are able to cope with seasonally varying climatic stressors through developmental as well as adult acclimation during their life time (Denlinger and Lee, 2010). However, lesser attention has been paid to acclimatization of tropical ectothermic organisms (Whitman and Ananthakrishnan, 2009). Tropical drosophilids encounter significant seasonally varying low vs high relative humidity conditions while thermal changes are quite limited i.e. 24 to 30 °C (www.tropmet.res.in).

In tropical regions, seasonal changes in humidity conditions play a major role in affecting morphological, physiological and life history traits (Tauber et al., 1986; Tauber et al., 1998). For example, role of humidity has been demonstrated for dry season induced diapause in some tropical insects (Pires et al., 2000; Seymour and Jones, 2000); increased desiccation resistance due to low humidity developmental acclimation in *D. leontia* (Parkash and Ranga, 2014); and due to effect of humidity acclimation on heat resistance in adult flies of *D. simulans* (Bubliy et al., 2013). Further, another study on tropical *D. jambulina* has shown effect of low vs high humidity acclimation on mating related traits of darker and lighter morphs consistent with melanism-desiccation hypothesis (Parkash et al., 2009). This study has shown that the frequencies of melanic and non-melanic morphs of *D. jambulina* are driven by humidity changes and not due to thermal conditions (Parkash et al., 2009). However, plastic changes induced by multiple stressors for tropical insect taxa of wet – dry seasons have received less attention (Tauber et al., 1998; Chown et al., 2011).

In Southeast Asia, changes in relative humidity associated with reduced patterns of rainfall due to El nino and climate warming are likely to increase drier conditions affecting survival of tropical insect taxa (www.skymetweather.com; www.imd.gov.in). Therefore, assessment of acclimatization potential of stenothermal tropical drosophilids reared under wet – dry conditions can help in understanding species specific stress resistance potential to multiple stressors and likely consequences on their fitness and survival (Hoffmann, 2010).

During their life time, ectothermic organisms (freeze tolerant, freeze avoiding or chill susceptible) undergo stressor induced plastic changes in stress resistance traits as well as metabolic fuels for survival under harsh climatic conditions (Denlinger and Lee, 2010; Sinclair et al., 2013). Analyses of cold induced plastic changes in metabolites have emphasized the role of different cryoprotective colligative solutes such as sugars, polyols and free amino acids (Overgaard et al., 2007; Michaud et al., 2008; Kostal et al., 2011b; Colinet et al., 2012). First, the role of exogenous trehalose to increase resistance to cold has been shown in *Belgica antarctica* (Benoit et al., 2009). Second, laboratory flies of *D. melanogaster* subjected to cold shock at -7 °C for 2-3 h; as well as strains resisting chilling injury at 0 °C for 30 to 60 h, revealed 3 to 6 fold increase in the level of proline (Misener et al., 2001). Accumulation of proline in response to cold stress has been demonstrated in the larvae of *D. melanogaster* (Kostal et al., 2011b); in *Chymomyza costata* larvae (Kostal et al., 2011a).Third, In the gall fly *Eurosta solidaginis,* cold induced energy metabolites include glycerol and sorbitol (Lee, 2010). Fourth, as compared to many studies on sugars and polyols, changes in free amino acids accumulated in response to different climatic stressors have been investigated in few insect taxa (Fields et al., 1998; Misener et al., 2001); and in a arthropod (Issartel et al., 2005). Thus, plastic responses to cold involve different physiological mechanisms based on different energy metabolites.

In tropical habitats, heat as well as drought stress co-occur and proline has been shown to play a role in thermoprotection of some plant taxa (Verbruggen and Hermans, 2008). The role of proline as an osmolyte to mitigate water stress was first reported in wilting perennial rye grass -*Lolium perenne* (Kemble and MacPherson, 1954) and subsequently in bacteria (Csonka and Hanson, 1991). Higher levels of proline have been observed in *Arabidopsis thaliana* (Liang et al., 2013) and in drought-tolerant rice varieties (Choudhary et al., 2005) but not in case of barley (Chen et al., 2007). Therefore, associations between high proline levels and drought tolerance have shown mixed results in plants and need further studies. Proline has been considered as a multifunctional amino acid due to its role as cryoprotectant and / or osmoprotectant and in mitigating oxidative stress in plants (Szabados and Savoure, 2009). However, a recent study has shown accumulation of proline due to drought stress in *D. immigrans* (Tamang et al., 2017). Thus, despite the abundant level of proline in insects, its physiological role(s) in diverse insect taxa need further studies.

In ectothermic organisms, survival under harsh environments depends upon maintenance of energetic homeostasis in metabolic fuels after possible perturbations due to climatic stressors (Benoit et al., 2009; Sinclair et al., 2013; Williams et al., 2014; Sinclair, 2015). In this respect, some studies have evidenced cold induced changes in metabolites through targeted ^1^H NMR based metabolomics (Overgaard et al., 2007; Michaud et al., 2008; Colinet et al., 2012). Based on metabolic profiling of temporal changes (during recovery), protective role of cold or heat hardening is evident from faster attaining of homeostasis of metabolites to normal level as compared to perturbations in metabolic pool in the control groups of *D. melanogaster* (Malmendal et al., 2006; Overgaard et al., 2007; Kostal et al., 2011b; Williams et al., 2014). However, efforts to find marker metabolites induced by cold or drought or heat stress have shown similarities as well as differences. For example, Antarctic midge *Belgica antarctica* has shown some overlap in the accumulation of glycogen and erythritol due to cold or drought acclimation while reduction in the level of serine was evident in response to cold, drought and heat stress (Michaud et al., 2008). In nature, acclimatization of ectothermic organisms involves developmental as well as adult acclimation. Therefore, NMR metabolic profiling of field acclimatized insects may further help in understanding stressor induced plastic changes in the metabolome. Further, there is need to examine energy metabolites mediated cross-protection to multiple stressors in insect taxa living under seasonally varying environments of tropical regions.

Temperature is considered as a major abiotic factor affecting geographical distribution and abundance levels of various insect taxa (Andrewartha and Birch, 1970; Angilletta, 2009). Narrow distribution patterns of tropical *Drosophila* species are limited by their low genetic potential to adapt to colder and drier habitats (Kellermann et al., 2009). Further, in context of climate warming, it has been argued that tropical drosophilids might be vulnerable due to expected higher aridity conditions (Hoffmann, 2010; Chown et al., 2011). Despite the fact that plastic changes in stress resistance traits in the generalist species, *D. melanogaster* are higher as compared with genetic differences, similar studies have not been carried out for tropical *Drosophila* species (Hoffmann et al., 2005). On the Indian subcontinent, there are contrasting seasonal patterns of ambient relative humidity (80 ± 5% RH during monsoon but 40 ± 6 % RH during autumn). Therefore, assessment of season specific humidity acclimation on induced stress resistance as well as their cross-tolerance effects can help in understanding adaptive potential of tropical *D. ananassae.*

In the present work, we assessed season-specific as well as sex-specific plastic changes in stress related traits as well as changes in the levels of three energy metabolites in the tropical *D. ananassae* which is characterized by low desiccation resistance. Wild - caught flies of *D. ananassae* from wet (monsoon) and dry (autumn) seasons were reared under season specific simulated growth conditions and flies were tested for basal as well as induced level of resistance to heat, desiccation and starvation resistance. For each stressor, we tested cross resistance for other two stressors. Further, we investigated patterns of changes (accumulation and / or utilization) for each of the three metabolic fuels due to plastic changes. For control as well as acclimated flies, changes in energy budget were also estimated. The rates of change in the levels of three energy metabolites were assessed in three replicates of multiple groups of flies subjected to different time durations of heat hardening, desiccation or starvation acclimation. Thus, we aim to find stressor induced plastic changes for possible maintenance of energetic homeostasis in the tropical *D. ananassae* from wet or dry seasons.

## MATERIAL AND METHODS

### Collection and Cultures

Wild-caught *Drosophila* species were collected during two seasons wet or rainy (July and August) and dry or autumn (Mid September to mid November) from five local sites but each one separated by ∼ 3 to 4 km in the university town Rohtak (Latitude 28.08 °N, Altitude 220m) in the year 2015. The flies were collected during one week in the mid of rainy or autumn season. Based on our past collection records, the relative abundance of *Drosophila ananassae* is ∼30% in rainy season and ∼20% in the autumn. Therefore, we collected 730 flies in rainy season and 416 flies in autumn season which included different *Drosophila* species. However in the laboratory, assortment of wild-caught flies provided 192 flies of *D. ananassae* in rainy season and 136 in autumn season. For each season, flies were used to set up three replicate populations in 300 ml culture bottles, each with 40 pairs of *D. ananassae.* Further, adult flies of each bottle were allowed to oviposit on cornmeal-yeast-agar medium in four culture bottles in order to maximize the number of laboratory raised flies. The wet season wild-caught flies were reared under wet season specific simulated condition (26±1°C and 78±2% RH). The resulting adult flies of G1 and G2 were used for assessment of basal, acclimated, cross-tolerance effects of three stressors (heat, drought and starvation) along with simultaneous analysis of control or unacclimated flies. For wet season, one week old flies of G_1_ and G_2_ generations were analyzed for changes in three stress related traits as well as energy metabolites (body lipids, proline and trehalose). Thus, all experiments on wet season flies were completed before the onset of autumn season. Likewise, for autumn or dry season, collection of wild flies, setting up of mass populations in triplicate and rearing under dry condition (25±1°C; 40±2% RH) were conducted during the autumn season. For dry season flies, 1stress resistance traits (heat knockdown, desiccation or starvation resistance) as well as energy metabolites (trehalose, proline and total body lipids contents), were assessed in three replicates of thirty flies. Control experiments were run simultaneously.

### Stress resistance assays

**(a)** Heat knockdown time was measured in three replicates of thirty flies of both the seasons. Individual males and females were placed into 5mL glass vials submerged into a water bath at a constant temperature of 37 °C. Flies were scored for knockdown time (in minutes and seconds).

**(b)** Desiccation resistance was measured as the time to dehydration effect under dry air (∼8% RH) in flies of both the sexes of wet or dry season. Each vial contained 2 g of silica gel at the bottom overlain with a foam disc to avoid contact with flies. We placed ten flies in such plastic vials (40 x 100 mm) in which open end was covered with muslin cloth. Finally, such vials were kept in the desiccator chamber (Secador electronic desiccator cabinet; www.tarsons.in) which maintained ∼8% relative humidity. Number of immobile flies was counted after hourly intervals; and LT100 values were recorded.

**(c)** For three replicates of thirty flies, starvation resistance was measured as survival time till death under humid conditions (∼90% RH) but without food. Ten adult flies were placed in a dry plastic vial which contained foam sponge impregnated with 8 ml of water + 2 mg sodium benzoate (to prevent any bacterial growth). The mortality time was recorded twice a day till all flies died from starvation.

### Assessment of direct and cross-tolerance effects

Direct as well as cross-tolerance effects were assessed for each stressor, (heat or drought or starvation) in *D. ananassae* flies of wet or dry seasons. For each stressor, different groups (three replicates of thirty flies each) of acclimated flies were tested for changes in other two stress related traits. Thus, for three stressors (heat or desiccation or starvation), we tested nine acclimation-by-test combinations (i.e. three direct effects + six cross tolerance effects). For analysis of cross-tolerance, acclimated/hardened fly groups were tested for changes (increase/decrease/no effect) in the level of each stress resistance trait. Therefore, for testing direct and cross tolerance due to (i) heat hardening, three groups of thirty flies of both the sexes were subjected to 2 h heat stress followed by 2 h recovery period on nutrient agar medium; and thereafter flies were analyzed for change in heat knockdown, desiccation or starvation resistance. (ii) Desiccation acclimation was given to flies for 4 h followed by 4h recovery period; and these flies were analyzed for heat knockdown, desiccation and starvation resistance. (iii) Flies were exposed to 20 h for starvation acclimation followed by 15h recovery period; followed by testing for changes in heat knockdown, starvation and desiccation resistance.

### Estimation of energy metabolites

**(a)** Body lipid content was estimated on G1 or G2 flies (three replicates of thirty flies of each season and sex) reared under wet or dry season specific conditions. For lipid content individual fly was dried in 2 ml Eppendorf tubes (http://www.tarsons.in) at 60 °C for 48 h and then weighed on Sartorius microbalance (Model-CPA26P; 0.001 mg precision; http://www.sartorious.com). Thereafter, 1.5 ml di-ethyl ether was added in each Eppendorf tube and kept for 24 h under continuous shaking (200 rpm) at 37 °C. Finally, the solvent was removed and individuals were again dried at 60 °C for 24 h and reweighed. Lipid content was calculated per individual by subtracting the lipid-free dry mass from initial dry mass per fly.

**(b)** For trehalose content estimation, each of the three replicates of thirty flies of each season and sex were homogenized in a homogenizer (Labsonic@ M; http://www.sartorious.com) with 300 μl Na2CO3 and incubated at 95 °C for 2 h to denature proteins. An aqueous solution of 150 μl acetic acid (1 M) and 600 μl sodium acetate (0.2 M) was mixed with the homogenate. Thereafter, the homogenate was centrifuged (Fresco 21, Thermo-Fisher Scientific, Pittsburgh, USA) at 12,000 rpm (9660 ×g) for 10 min. For trehalose estimation, aliquots (200 μl) were placed in two different tubes; one was taken as a blank whereas the other was digested with trehalase at 37 °C using the Megazyme trehalose assay kit (K-Treh 10/10, http://www.megazyme.com). In this assay, released D-glucose was phosphorylated by hexokinase and ATP to glucose-6-phosphate and ADP, which was further coupled with glucose-6-phosphate dehydrogenase and resulted in the reduction of nicotinamide adenine dinucleotide (NAD). The absorbance by NADH was measured at 630 nm (UV-2450-VIS, Shimadzu Scientific Instruments, Columbia, USA). The pre-existing glucose level in the sample was determined in a control reaction lacking trehalase and subtracted from total glucose concentration.

**(c)** Proline content was estimated in each of the three replicates of thirty flies of each season and sex. Proline concentrations in fly homogenates were determined by the modified method after Bergman and Loxley (1970). In this assay, interference from primary amino acids gets eliminated by nitrous acid treatment and the excess nitrous acid is removed by heating with ammonium chloride followed by hydrochloric acid. Interfering materials are also removed by absorption to the protein-sulphosalicylic acid complex.

Thirty adult flies were homogenized in 3 ml of sulphosalicylic acid. Following centrifugation, 50 μl of the homogenate was added to 15 μl of freshly prepared 1.25 M sodium nitrite solution and content were mixed and kept at room temperature for 20min. Further, 15 μl of 1.25Mammoniumchloride solution added and content were mixed followed by addition of 60 μl of concentrated hydrochloric acid. The content were mixed and heated in a boiling water bath for 20 min. The tubes were cooled and 60 μl of 10 N sodium hydroxide was added. To the resulting solution, we added 200 μl glacial acetic acid and 200 μl of ninhydrin solution in each capped tube. The solutions were mixed and incubated for 60 min. at 100 °C. Following incubation, the samples were extracted with toluene, and absorbance of the aqueous phase was quantified spectro-photometrically at 520 nm and the amount of proline was estimated in reference to a standard curve.

### Analysis of rate of change in energy metabolites after hardening/acclimation pretreatments

We conducted independent experiments for each stressor to find change in the rate of accumulation or utilization of three energy metabolites (trehalose, proline and body lipids). Such changes were assessed in three replicates of thirty flies of each season as well as sex as a function of different time durations of hardening/ acclimation by a stressor. Independent groups of flies were (a) heat hardened for 1h, 2h, 3h, 4h and 5h at 34°C with 2h recovery;(b) desiccation acclimation for1h, 2h, 3h, 4h and 5h at 8% relative humidity with 4h recovery; (c) starvation acclimation for 10h,15h, 20h, 25h and 30h followed by 15h recovery; and these respective flies were tested for changes in the level of each of three energy metabolites.

### Assessment of energy metabolites mediated cross-protection

For acclimated as well as non-acclimated (control flies), changes in three energy metabolites (trehalose, body lipids and proline content) due to each stressor were measured in three replicates each of thirty flies. This was done for flies reared under season-specific wet or dry conditions. For such analysis, we used data on flies after heat hardening (2 h heat stress followed by 2 h recovery); (ii) desiccation (5 h acclimation followed by 4 h recovery); (iii) starvation (25 h starvation acclimation followed by 15h recovery). Finally, percent change in accumulation or utilization of three energy metabolites due each stressor were calculated to find possible cross-protection between multiple stressors.

### Treatment and analysis of data

Data for heat knockdown time at 37°C (in minutes and seconds) were recorded on individual male and female flies of *D. ananassae* reared under wet or dry season specific simulated condition. For other two stressors (desiccation or starvation), we recorded survival mortality of 10 flies per vial as a function of shorter (at hourly) for desiccation; and twice daily (8am and 8pm) for starvation resistance till all the flies died. For each season and sex, (three replicates of thirty flies), data on basal level (or control) and hardened/acclimated flies were represented as mean ± s.e.m. (Table 1) while effects of treatments and sex were calculated on the basis of ANOVA (supplementary tables). Seasonal differences in stress related traits were compared with ‘t’test as well as in terms of fold differences (Table 1).

**Table 1.** Data on seasonal differences in basal levels of stress resistance to heat, or desiccation or starvation stress and energy metabolites (trehalose, proline and body lipids) of *D. ananassae* flies grown under season specific simulated conditions (rainy or autumn). For each trait, data are mean ± s.e.m for three replicates of thirty flies (n = 90).

|  | Traits | Sex | Wet season | Dry season | Fold | *t*-test |
| --- | --- | --- | --- | --- | --- | --- |
| (a) | Heat Knock-down (min.) | F | 109 ± 2.32 | 98 ± 3.12 | 1.11 | 12.78*** |
| | | M | 100 ± 4.13 | 86 ± 2.20 | 1.16 | 23.66*** |
| (b) | Desiccation resistance (h) | F | 12 ± 0.56 | 16 ± 1.02 | 1.33 | 5.09*** |
| | | M | 10 ± 0.37 | 14 ± 0.78 | 1.40 | 4.48*** |
| (c) | Starvation resistance (h) | F | 70 ± 2.13 | 41 ± 3.42 | 1.70 | 15.25*** |
| | | M | 53 ± 1.09 | 36 ± 3.02 | 1.47 | 32.22*** |
| (d) | Trehalose ($\mu\text{g mg}^{-1} \text{ fly}^{-1}$) | F | 11.20 ± 0.25 | 24.74 ± 0.44 | 2.20 | 18.72*** |
| | | M | 17.71 ± 0.13 | 22.31 ± 0.35 | 1.26 | 12.20*** |
| (e) | Proline ($\mu\text{g mg}^{-1} \text{ fly}^{-1}$) | F | 13.48 ± 0.11 | 48.55 ± 1.06 | 3.60 | 27.12*** |
| | | M | 18.19 ± 0.29 | 52.48 ± 1.48 | 2.89 | 45.40*** |
| (f) | Body lipids ($\mu\text{g mg}^{-1} \text{ fly}^{-1}$) | F | 190 ± 3.55 | 123 ± 2.56 | 1.54 | 56.30*** |
| | | M | 140 ± 3.02 | 108 ± 2.26 | 1.30 | 43.21*** |

\*\*\* $P < 0.001$

The data on assays for three stressors (heat, drought or starvation) were used for calculating absolute acclimation/hardening capacity -AAC (i.e. difference in trait values between acclimated flies - control flies) following Kellett et al (2005). Further, we also calculated relative acclimation capacity - RAC (i.e. absolute acclimation capacity divided by control value of unacclimated flies). For all the three stress related traits, we represented AAC in the form of bars while RAC values were depicted on the top of each bar. Illustrations depicted simultaneous comparison of direct acclimation effect as well as cross tolerance effects for male and female flies of wet or dry season (Fig 1 to 3). For heat resistance, correlation between direct heat hardening effect and cross-tolerance effects due to desiccation acclimation was represented (with Box-whisker) for flies of two seasons as well as sexes. Further, we represented relationship between heat hardening effect on starvation as well as on heat knockdown (Fig. 4).

**Fig. 1.**
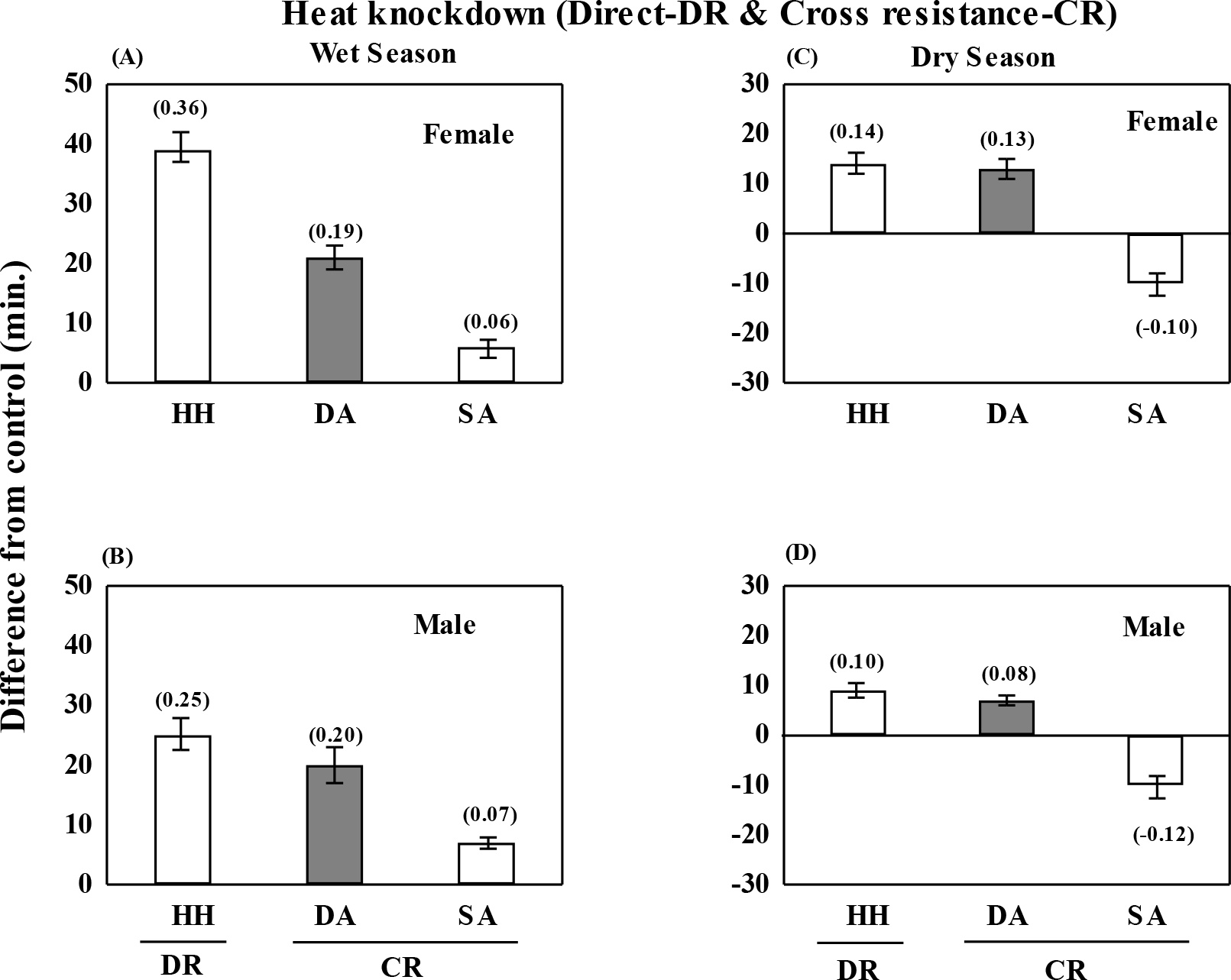
Plastic changes (acclimated trait value – control) in heat knockdown (minutes) as a consequence of heat hardening (direct effect) and due to cross tolerance effects of desiccation - DA or starvation – SA in *D. ananassae* of wet or dry season. Relative acclimation capacity (RAC) value is shown on top of each bar.

**Fig. 2.**
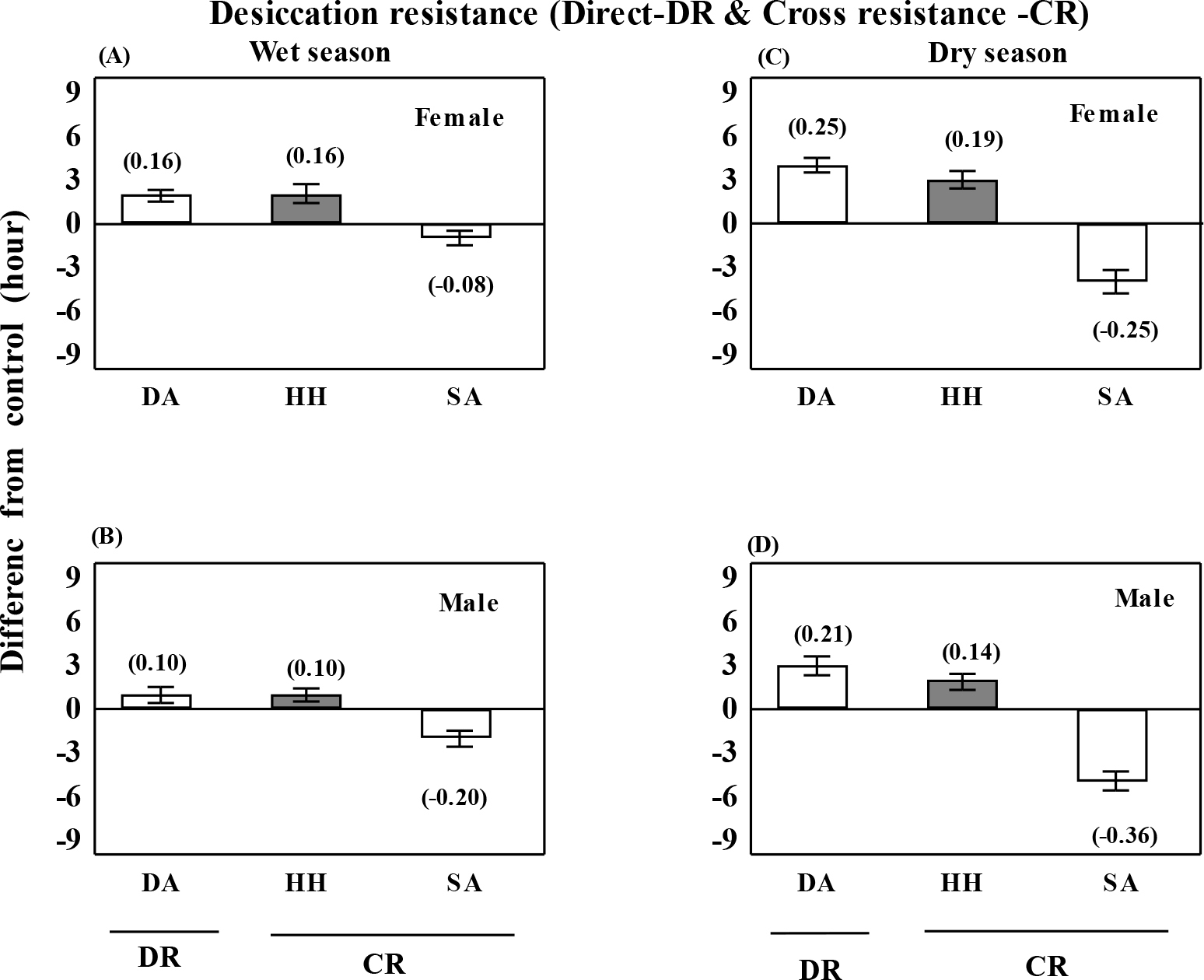
Wet or dry season as well as sex specific changes in desiccation resistance (h) as a consequence of desiccation acclimation (direct effect) and due to cross tolerance effects of heat hardened - HH or starvation acclimated -SA in *D. ananassae* of wet or dry season.

**Fig. 3.**
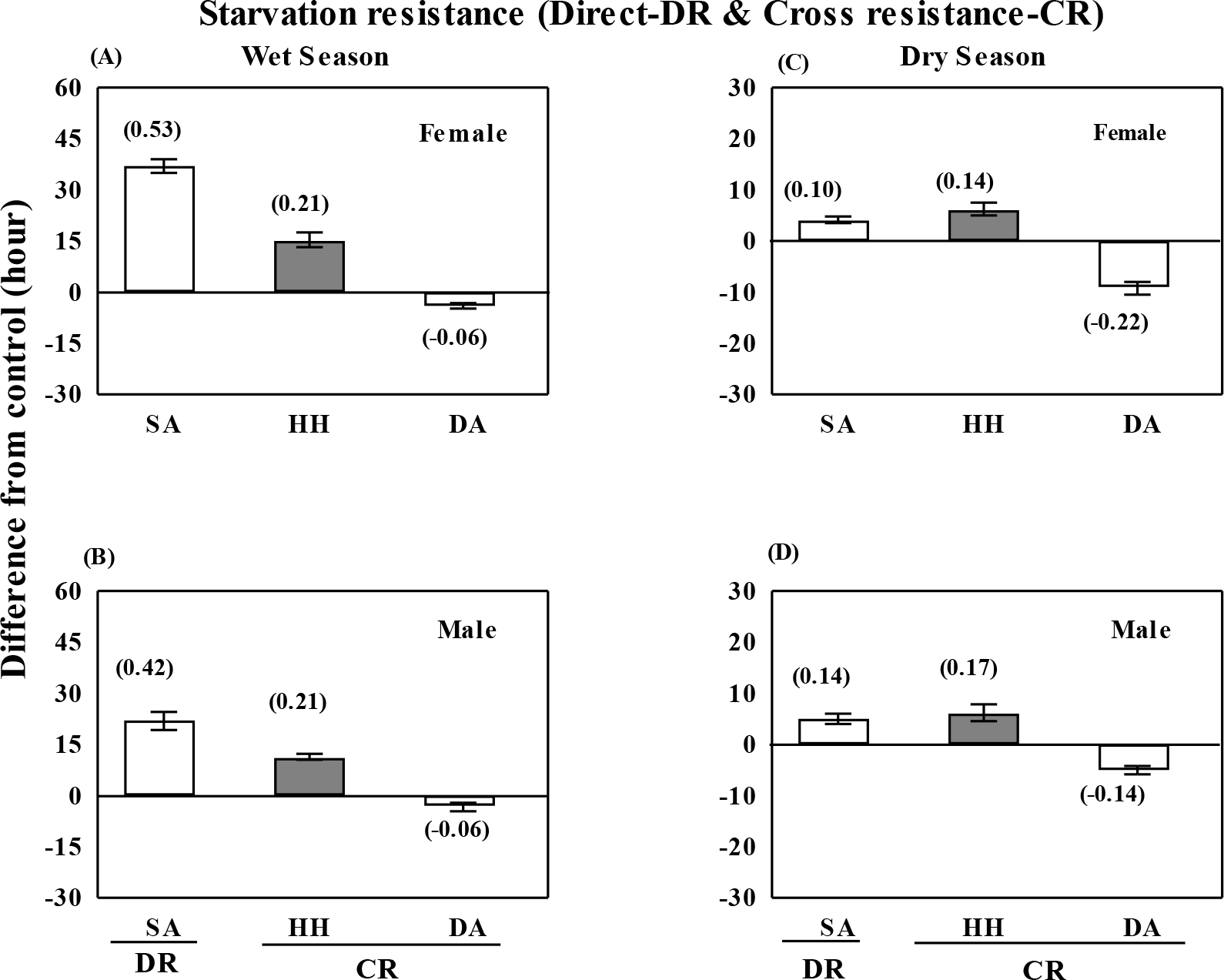
Comparison of plastic changes in starvation resistance (h) due to starvation acclimation (direct effect) and cross tolerance effects of heat hardened - HH or desiccation acclimated – DA in *D. ananassae* of wet or dry season.

**Fig. 4.**
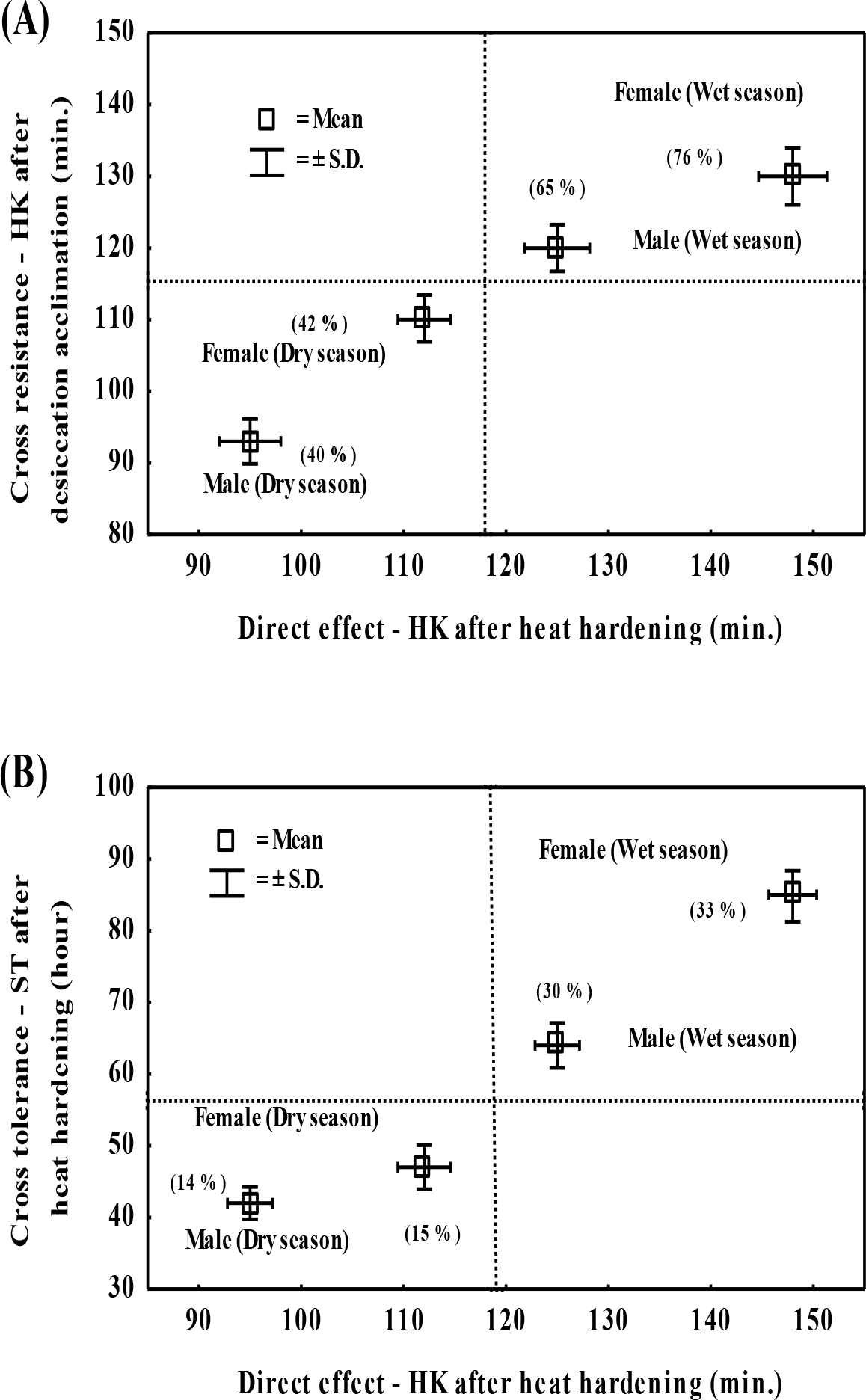
(A) Relationship between plastic changes in heat knockdown due to direct effect of heat hardening (x-axis) versus cross tolerance effect due to desiccation acclimation (y-axis). (B) Correlation between plastic changes in heat knockdown as well as starvation resistance due to heat hardening in *D. ananassae* of wet or dry season. Percent changes in energy budget per fly due to plastic changes are indicated in parentheses.

In accordance with the objectives of this study, data on sum of three energy metabolites (ug mg^-1^ fly^-1^) in control flies were compared with heat hardened or flies acclimated to desiccation and starvation; and percent changes (+ or -) were calculated to compare acclimation effects across seasons and sexes (Table 2).

**Table 2.** Seasonal changes in the sum of energy budget per fly (for trehalose, proline and body lipids) due to direct-adult acclimation effects of heat hardening (2 h), desiccation acclimation (5 h) and starvation acclimation (25 h) of wet or dry season flies of *D. ananassae.* For each trait, data are shown as J mg^-1^ fly^-1^ along with percent change in parenthesis as compared with control (non acclimated flies).

|  | Sum of energy budget (females) |  | Sum of energy budget (males) |  |
| --- | --- | --- | --- | --- |
|  | Wet season flies | Dry season flies | Wet season flies | Dry season flies |
| HA | 13.66 ± 0.91 (+ 72.85) | 7.94 ± 0.33 (+ 29.63) | 9.91 ± 0.98 (+ 61.44) | 7.09 ± 0.15 (+ 27.17) |
| DA | 8.17 ± 0.32 (+ 3.35) | 6.85 ± 0.29 (+ 11.72) | 6.37 ± 0.37 (+ 3.73) | 6.25 ± 0.24 (+ 12.35) |
| SA | 4.76 ± 0.16 (- 39.73) | 5.23 ± 0.11 (- 14.66) | 4.18 ± 0.28 (- 31.92) | 4.82 ± 0.36 (- 13.41) |
| Cont | 7.91 ± 0.24 | 6.13 ± 0.22 | 6.14 ± 0.33 | 5.57 ± 0.28 |

Heat hardening = HA; Desiccation acclimation = DA; Starvation acclimation = SA; Cont = Control
Conversion factors include 17.6 J mg<sup>-1</sup> for trehalose, 39.3 J mg<sup>-1</sup> for lipids, 17.8 J mg<sup>-1</sup> for proline (Schmidt-Nielsen. 1990).

Data obtained from independent sets of experiments on rate of metabolite change as a consequence of different durations (1h, 2h, 3h, 4h or 5h) for heat hardening or desiccation acclimation; were subjected to regression analysis for calculation of regression slope values (Table 3); and seasonal differences in slope values were compared with ‘t’ test. Finally, season specific differences in the accumulation and utilization (calculated in terms of percent increase or decrease) of three energy metabolites (body lipids, trehalose and proline) due to either heat hardening or acclimation to desiccation or starvation were schematically represented to highlight possible energy metabolite mediated cross-protection in wet or dry season female flies (Fig. 5). For stressor acclimated/hardened flies, the energy content (trehalose, proline and body lipids) was calculated using standard conversion factors (Schmidt-Nielsen, 1990). The amount of each energy metabolite was multiplied by conversion factor i.e. for trehalose (17.6 Jmg^-1^); for body lipid (39.3 Jmg^-1^); and proline (17.8 Jmg^-1^). Statistical calculations and illustrations were made with the help of Statistica 5.0 as well as Statistica 7.

**Table 3.** Rate of metabolite change (regression slope values) as a function of different durations of heat hardening or desiccation acclimation (1h or 2h or 3h or 4h and 5h); and starvation acclimation (10h or 15h or 20h or 25h and 30 h) for Wet and Dry season flies of *D. ananassae.*

| Rate of change in each metabolite under heat, desiccation or starvation stress |  |  |  |  |
| --- | --- | --- | --- | --- |
|  | Sex | Wet season | Dry season | *t*-test |
| **(a) Heat hardening ($\mu\text{g min}^{-1}$)** |  |  |  |  |
| Trehalose | F | -0.07 | -0.03 | 0.011** |
|  | M | -0.12 | -0.02 | 0.016*** |
| Proline | F | ----- | -0.20 | ----- |
|  | M | ----- | -0.23 | ----- |
| Body lipids | F | +1.25 | +0.49 | 0.020*** |
|  | M | +0.85 | +0.43 | 0.036*** |
| **(b) Desiccation acclimation ($\mu\text{g h}^{-1}$)** |  |  |  |  |
| Trehalose | F | +2.14 | +1.14 | 1.03*** |
|  | M | +3.02 | +0.96 | 0.98*** |
| Proline | F | ----- | +4.86 | ----- |
|  | M | ----- | +4.79 | ----- |
| **(c) Starvation acclimation ($\mu\text{g h}^{-1}$)** |  |  |  |  |
| Body lipids | F | -3.20 | -0.89 | 1.01*** |
|  | M | -2.00 | -0.80 | 0.94*** |

Slope values represent rate of change of metabolites as a function of time. \*\* $P < 0.01$ , \*\*\* $P < 0.001$

**Fig. 5.**
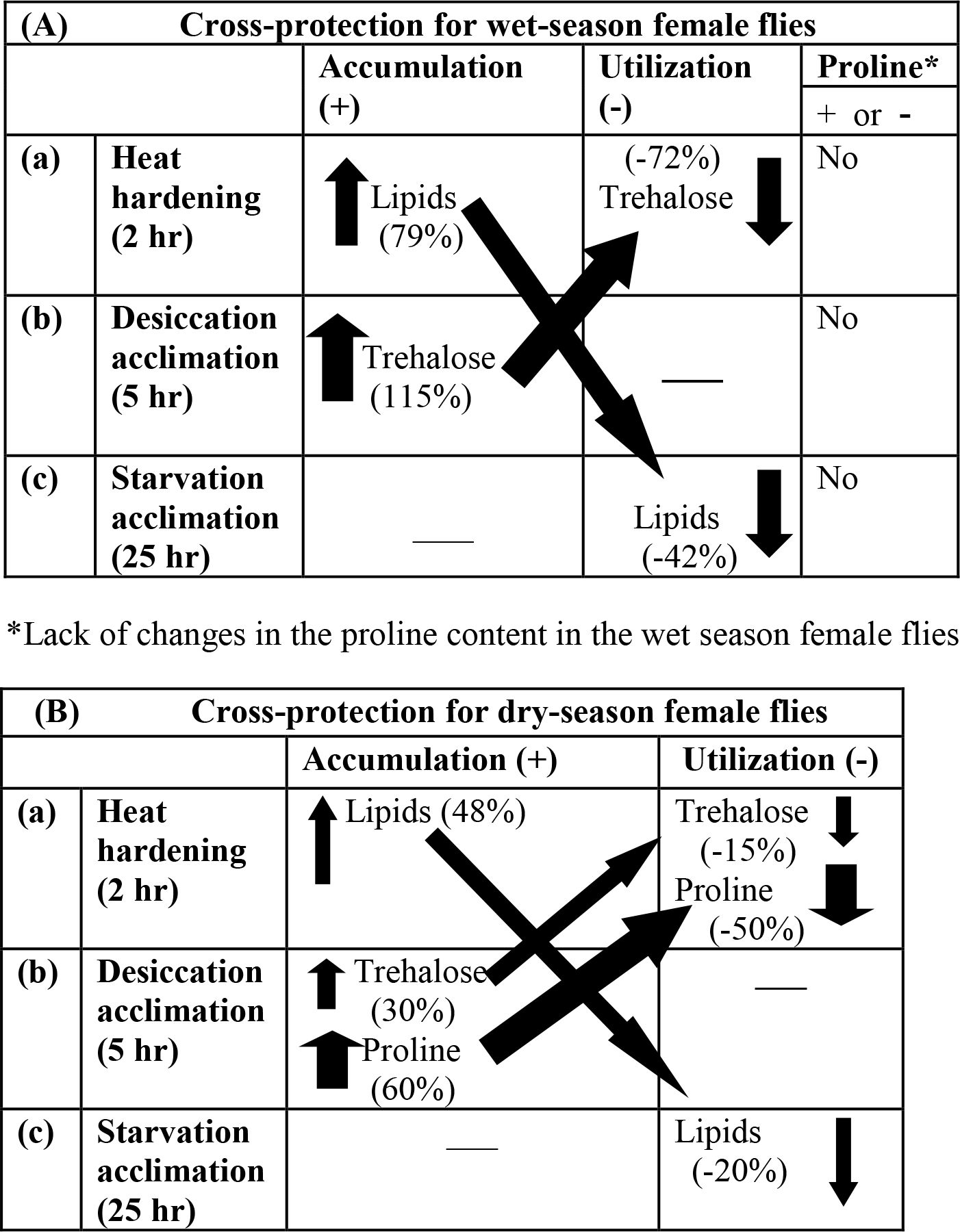
Schematic representation of stressor induced (heat or drought or starvation) plastic changes (accumulation and utilization) of three energy metabolites (trehalose, proline and body lipids) in *D. ananassae* female flies of wet (A) or dry (B) season. Slanted arrows depict cross-protection effects.

## RESULTS

### Seasonal differences in stress resistance and energy metabolites

Data on seasonal differences (wet vs dry) for six traits of *D. ananassae* flies reared under season specific simulated conditions are shown in Table 1. For heat knockdown, starvation resistance and body lipids, wet season flies revealed significantly higher trait values as compared with dry season flies (Table 1). For dry season flies, desiccation resistance and the amount of trehalose and proline were significantly higher. For all the traits, season specific differences were significant for both the sexes (*p* < 0.001; Table 1).

### Plastic changes in heat knockdown of wet vs dry season flies

For heat knockdown, data on absolute hardening capacity (acclimated - control values) due to direct hardening as well as cross tolerance effects due to desiccation or starvation acclimation are illustrated in Fig. 1. The wet season flies (both sexes) showed significant increase in heat knockdown due to heat hardening as well as cross resistance effect of desiccation (Fig. 1A,B). However, starvation acclimated flies of wet season showed lesser increase in heat knockdown (Fig. 1A,B). In contrast, in dry season flies, heat knockdown duration decreased as a consequences of cross tolerance effect due to starvation acclimation (Fig. 1C). However, plastic changes in heat knockdown of dry season flies due to direct heat hardening and cross tolerance effect after desiccation acclimation were 50 to 60% lower (Fig. 1C,D). Thus, direct effect due to heat hardening as well as cross tolerance effects significantly improved heat resistance of wet season flies as compared with dry season flies (Fig. 1).

### Season specific plastic changes in desiccation resistance

Wet season flies exhibited lower acclimation effects due to desiccation acclimation as well as cross tolerance effect of heat hardened flies (Fig. 2A,B) as compared with dry season flies (Fig. 2C,D). There was a trade-off between desiccation and starvation resistance. We observed two fold reduction in desiccation resistance of starvation acclimated flies of dry season as compared with wet season.

### Seasonal plastic changes in starvation resistance

For starvation resistance, wet season flies exhibited significantly higher effect of direct starvation acclimation as well as cross-tolerance due to heat hardening as compared with dry season flies (Fig. 3). In case of desiccation acclimated flies, there was greater reduction in starvation resistance of dry season flies as compared with wet season flies.

### Relationship between heat hardening effects with other traits

*D. ananassae* flies are acclimatized to multiple stressors in its natural habitats across season. For seasonal changes in heat knockdown, we found positive relationship between direct heat hardening and cross tolerance due to desiccation acclimation (Fig. 4A). Thus, significant increase in heat resistance of wet season flies results due to plastic effects of heat as well as drought (Fig. 4A). Inter-related plastic changes between resistance to heat and starvation as a consequence of heat hardening are shown in Fig. 4B. Therefore, heat hardening of wet season flies resulted in higher resistance to both heat and starvation while such effects were lower for dry season flies.

### Assessment of relative hardening / acclimation capacity

Results of relative acclimation capacity (RAC) for direct as well as cross-tolerance plastic effects for three stressors are shown in Fig. 1-3. In wet season flies, direct acclimation effects were maximum for starvation acclimation followed by heat hardening. However, direct effect of desiccation acclimation was higher for dry season flies as compared with wet season flies. RAC values for cross-tolerance effects (SA on HK); (DA on HK) and (DA on HH) were quite similar i.e. RAC = ∼0.20 (Fig. 1-3). These observations suggest that RAC values for direct acclimation / hardening are stressor specific while cross tolerance effects could be conserved for a species. This assumption needs further verification for its generality among *Drosophila* species. Further, we observed a trade-off between RAC values of cross-tolerance effects of DA vs SA but such effects were significantly higher for dry season flies (Fig. 2 & 3). Finally, cross-tolerance effect of SA on heat knockdown was positive in wet season flies but there was a trade off (SA on HK) in dry season flies (Fig. 1). We also found sex-specific minor differences in RAC for stress related traits of wet or dry season flies (Fig. 1-3).

### Season specific plastic changes in energy budget

Energy budget based on three energy metabolites (trehalose, proline and body lipids per fly) in control and heat hardened or flies acclimated to desiccation or starvation are shown in Table 2. There is significant increase (∼ 73% in females and ∼ 62% in male) in the energy budget of wet season flies due to heat hardening (Table 2) while such changes in the energy budget of dry season flies are sixty percent lower. In contrast, there is ∼ 12% gain in energy budget due to desiccation acclimation of dry season flies as compared with ∼ 4% in the wet season flies. Further, starvation acclimated flies of wet season consumed about 40 percent of energy budget per fly as compared with ∼ 14% in dry season flies. These observations show plastic changes in energy budget are stressor as well as season specific while both the sexes reveal similar trends.

### Cross-protection between accumulation and utilization of energy metabolites

In *D. ananassae* flies reared under wet or dry conditions, we found significant differences in the stressor specific levels of accumulation and utilization of trehalose, proline and body lipids (Fig. 5). This schematic diagram shows that body lipids increase due to heat hardening i.e. ∼ 79% in wet season flies as compared with 48% for dry season flies. There is cross-protection between accumulation of body lipids due to heat hardening and utilization under starvation. In contrast, proline was elicited by desiccation acclimated flies of dry season only. However, desiccation acclimated wet season flies accumulated 115% more trehalose than control flies. Further, heat hardening of wet season flies utilized 50% of the accumulated trehalose. For dry season flies, desiccation induced plastic changes accumulated only 30% more trehalose but 60% more amount of proline which were partially utilized under heat hardening. Thus, plastic changes in three energy metabolites (trehalose, proline and body lipids) are consistent with cross-protection between three stressors (heat, drought and starvation).

### Stressor specific rates of change in energy metabolites

We found significant seasonal differences in stressor specific (heat, drought or starvation) changes (accumulation or utilization) in the levels of each of the three energy metabolites as a function of different durations of hardening / acclimation of wet or dry season flies (Table 3). For body lipids and trehalose, the rates of change were significantly higher for wet season flies as compared with dry season flies. Sex-specific differences in the rates of metabolite change were observed only in the wet season flies (Table 3). Further, changes in the level of proline were found only in dry season flies.

### Results of ANOVA on stress related traits

In supplementary Table S1, we have shown results of ANOVA for three stress related traits (heat, drought or starvation) with respect to three variables (control vs acclimated; sexes and seasons) in *D. ananassae* flies reared under wet or dry season specific conditions. For all the three stress related traits, we found highly significant differences (*p* <0.001) across sexes, seasons and due to acclimation as well as due to their respective interactions (Table S1). Similarly the results of ANOVA on three energy metabolites are shown in supplementary table S2 which also showed significant differences (*p* < 0.001) for all the variables.

## DISCUSSION

We observed humidity driven significant plastic changes in the stress related traits (heat knockdown, and resistance to desiccation or starvation); and three energy metabolites (proline, trehalose and body lipids) in *D. ananassae* flies reared under wet or dry season specific simulated conditions. Basal levels of stress resistance and energy metabolites differ significantly due to developmental acclimation. The effects of heat hardening as well as desiccation acclimation significantly improved heat knockdown in wet season flies. Further, heat hardening also increased starvation resistance in wet season flies. However, dry season flies showed higher levels of proline as well as desiccation resistance but a lower amount of trehalose. Therefore, proline can be considered as a marker metabolite because accumulation of proline was evident only in drought acclimated flies of dry season. Stressor specific changes in energy budget per fly support cross-protection between heat hardening and desiccation or starvation resistance. Thus, seasonal differences in relative humidity (wet or dry) induced cross-protective plastic changes in three energy metabolites are consistent with energetic homeostasis and for coping season specific stressful environments.

### Seasonal differences in cross-tolerance effects

Previous studies on cross-tolerance to multiple stressors did not consider impact of both developmental as well as adult acclimation effects of wet or dry conditions (Angilletta, 2009; Sinclair et al., 2013). In the present work, we examined such acclimation effects on tropical *D. ananassae.* In heat hardened flies (of wet or dry seasons), cross-tolerance effects on desiccation was higher for dry than wet season flies. However, such cross-tolerance effect on starvation tolerance was more in wet than dry season flies (Fig. 2,3). In contrast, in starvation acclimated flies cross-tolerance effects on heat resistance showed season specific contrasting differences i.e. a positive effect in wet season flies (Fig. 1A,B) but a negative effect in dry season flies (Fig. 1C,D). Therefore, cross-tolerance effects of starvation acclimated flies on heat knockdown differ across seasons. Further based on genetic effects, a previous study has shown lack of trade-off between resistance to heat and other stress related traits in *D. melanogaster* (Williams et. al., 2012). Thus, genetic and plastic effects of multiple stressors seem to differ in impacting heat tolerance but this assumption needs further analysis. In *D. ananassae,* cross-tolerance effects in two cases (a) plastic response of starvation acclimation on desiccation resistance (Fig. 2); (b) effect of desiccation acclimation on starvation resistance (Fig. 3) showed trade-off with greater effect in dry than wet season flies (Fig. 2,3).These observations are consistent with a trade-off between resistance to desiccation or starvation in geographical populations of some tropical drosophilids on the Indian subcontinents (Parkash and Munjal, 1999). Thus, we find similarity between plastic and genetic effects for desiccation vs starvation resistance in geographical as well as seasonal populations of tropical *Drosophila* species.

### Stressor specific relative acclimation capacity differ across wet-dry seasonal flies

Relative acclimation capacity (RAC) is a quantitative measure to compare stressor induced plastic changes across species as well as within species i.e. seasonal or geographical populations of ectothermic organisms (Kellett et al., 2005). The magnitude of RAC for heat knockdown has been found to be similar (RAC = 0.25) in interspecific comparisons of four species each of *melanogaster* species group and montium species group (Kellett et al., 2005). In contrast, in *D. melanogaster* flies reared at 18, 25 and 28 °C showed variable effects of thermal developmental acclimation on heat knockdown i.e. RAC ∼ 0.35 for 18 or 25 °C reared flies but no RAC effect for 28 °C flies (Cavicchi et al., 1995). Further, geographical populations of *D. melanogaster* from east coast of Australia revealed RAC values in the range of 0.13 to 0.58 for heat knockdown (Sgro et al., 2010). However, we are not aware of studies related to assessment of quantitative differences in stressor specific relative acclimation capacity in seasonally varying populations of diverse insect taxa. In the present work, we found significant season specific (wet or dry) as well as stressor specific differences in relative acclimation capacity (Fig. 1-3). For direct acclimation effects, RAC values were in the order of SA = 0.53 > HH = 0.36 > DA = 0.25. In contrast, cross-tolerance RAC values were quite similar (RAC = ∼ 0.20) for HH on SR, DA on HK (wet season flies) and HH on DR (dry season flies). The generality of such observations need further studies. Another interesting observation was cross-tolerance effect of starvation acclimation on heat knockdown i.e. a positive effect in wet season flies but a negative relationship (trade-off) in dry season flies. Thus, we found season specific and stressor specific differences in RAC effects in low vs high humidity reared flies of *D. ananassae.*

### Wet-dry seasonal flies differ in dehydration induced accumulation of osmoprotectants

Cold induced plastic changes in energy metabolites have revealed accumulation of trehalose as osmoprotectant (Shimada and Riihima, 1990; Benoit et al., 2009; Colinet et al., 2012) but not in *Sarcophaga crassipalpis* (Michaud and Denlinger, 2007). In the present work, drought induced accumulation of proline is associated with acclimatization of *D. ananassae* to drier conditions. *D. ananassae* flies (reared under dry season condition) showed significant accumulation of proline but a lower level of trehalose. However, *D. ananassae* flies reared under wet condition did not elicit proline at all despite accumulation of a significant level of trehalose (Fig. 5). Further, drought induced plastic changes in proline have been observed both for a cold adapted *D. immigrans* (Tamang et al., 2017) as well as warm adapted *Z. indianus* (Kalra et al., 2017). These studies suggest a possible role of proline as osmoprotectant in drosophilids reared under either colder and drier; or warmer and drier conditions. However, further studies are needed to support such observations.

### Cross-protection through inter-related plastic changes in metabolic fuels

In wild habitats, colder or warmer environments are coupled with wet or dry or possible starvation conditions; and insects are likely to evolve cross-protection mechanisms to multiple stressors. It is known that cold stress with coupled drought level is able to elicit multifunctional colligative solutes (sugars, polyols, proline as free amino acids). Possible inter-related modulatory changes (cross-protection) in the accumulation and utilization of metabolic fuels could favor energetic homeostasis. For example, a previous study has shown drought stress induced higher level of proline but exihibited its utilization under cold stress in winter population of *D. immigrans* (Tamang et al., 2017). In present work on tropical *D. ananassae,* we observed cross-protection between heat hardening and two coupled stressors i.e. drought or starvation. We found inter-related changes in three metabolic fuels both for wet or dry season flies of *D. ananassae.* Inter-relationship between heat hardening induced accumulation of body lipids and its consumption in starvation acclimated flies was evident for both the seasons but body lipid changes were 60 % higher for wet season flies as compared with dry season. Such observations are consistent with season specific differences in starvation resistance levels due to developmental and adult acclimation. However, inter-related metabolic changes due to heat hardening or desiccation acclimation involved accumulation and utilization of proline only in dry season flies. In contrast, we observed four fold higher (∼ 115 %) accumulation of trehalose in wet season flies as compared to dry season flies. Trehalose as osmoprotectant is known to counter the detrimental effects of drought stress on cellular membranes and cellular proteins. Thus, plastic changes in metabolic fuels due to multiple stressors seem to involve inter-related compensatory mechanisms to cope with the season specific wet or dry conditions.

Energetic changes in stressor induced and control groups of flies can also help in understanding cross-protection. Energy budget changes per fly (based on three metabolites) were significantly higher (∼ 70%) in wet season flies subjected to heat hardening but only (∼ 4%) due to desiccation acclimation while starvation acclimation consumed energy budget (∼ 40 %) as compared to control or unacclimated flies (Fig.5). Thus, wet season flies facing multiple stressors (heat or drought or starvation) are able to maintain its energy budget homeostasis. In case of dry season flies, energy budget changes revealed ∼ 30% increase due to heat hardening and 12 % with desiccation acclimation while starvation acclimation consumed 15 %, thereby showing a favorable energy balance under harsh dry conditions (Fig. 5). Thus, inter-related stressor induced changes in three energy metabolite seem consistent with possible energetic homeostasis.

### Thermoprotective effects of proline and trehalose

Some studies have examined detailed mechanisms for thermal protection of various cellular proteins and cell membranes by trehalose and / or proline supporting their ubiquitous role in stabilizing cellular components against detrimental effects of cold or heat (Kaushik and Bhatt, 2003; Yancey et al., 1982; Reina-Bueno et al., 2012; Liang et al., 2013). Both proline and / or trehalose provide cryoprotection in insects such as temperate *Chymomyza costata, Belgica Antarctica* and *D. melanogaster* adults as well as larvae (Kostal et al., 2011b; Benoit et al., 2009; Colinet et al., 2012; Kostal et al., 2011a) and also in different plant taxa (Szabados and Savoure, 2009). However, thermoprotective role of these two energy metabolites to heat stress have been less documented. First, metabolic profiling of heat hardened *D. melanogaster* adults showed elevated levels of alanine; and tyrosine (a precursor of stress hormones in insects) during recovery period (Malmendal et al., 2006). Since, alanine results due to proline oxidation during energy generation, utilization of proline can be argued during recovery of heat hardened *D. melanogaster* as reported by Malmendal and colleagues (2006). Second, association between heat and proline has been observed in beetle- *Alphitobius diaperinus* maintained at 28 °C because proline amount constituted 50% out of a pool of 16 amino acids (Renault et al., 2006). Third, thermoprotective role of proline as well as of trehalose has been suggested in some plant taxa (Verbruggen and Hermans, 2008; Szabados and Savoure, 2009); and in soil bacterium *Rhizobium etli* (Reina-Bueno et al., 2012). For *D. ananassae,* we observed partial utilization of proline when dry season flies were subjected to heat hardening (Fig. 5) but we did not examine possible elevation of alanine due to proline oxidation. Thus, further studies are needed to analyse osmo- as well as thermoprotective roles of proline in seasonally varying wild populations of different warm adapted drosophilids.

We observed complementary metabolic changes in accumulation and utilization of trehalose and proline in response to desiccation and heat stress in wet or dry season flies. Accumulation of proline occurred only in dry season flies in response to desiccation acclimation. We observed variable levels of trehalose in wet vs dry season flies. However, heat hardening resulted in partial utilization of trehalose in wet season flies but both proline and trehalose in dry season flies. Thus, proline could be a marker metabolite for dry season flies. Inter-related changes due to heat hardening and starvation acclimation involved accumulation and utilization of body lipids. Finally, we assessed energy budget changes per fly in control vs flies hardened / acclimated to heat or drought or starvation. For both the seasons (wet or dry) and sexes, heat hardening increased energy budget per fly due to build up of body lipids which were used during starvation. Thus, energy budget changes due to stressors resulted in cross-protection as well as maintenance of energetic homeostasis both under wet or dry climatic conditions. We may suggest that plastic induced changes in stress resistance traits and energy metabolites in *D. ananassae* are likely to counter future drier conditions expected due to climate change.

## Author contributions

A.P. and C.L. did laboratory as well as data analysis; and R.P. wrote the MS.

## References

Andrewartha, H.G. and Birch, L.C. (1970). The distribution and abundance of animals. Chicago University Press.

Angilletta, M.J. (2009). Thermal adaptation: A Theoretical and Empirical Synthesis. Oxford,UK: Oxford University Press.

Benoit, J.B., Lopez-martinez, G., Elnitsky, M.A., Lee, R.E. and Denlinger, D.L. (2009). Dehydration-induced cross tolerance of Belgica antarctica larvae to cold and heat is facilitated by trehalose accumulation. Comp. Biochem. Physiol.A. 152, 518-523.

Bergman, I. and Loxley, R.(1970). Improved spectrophotometric method for determination of proline in tissue hydrolysates. Anal. Chem. 42, 702-706.

Bubliy, O.A., Kristensen, T.N. and Loeschcke, V.(2013). Stress-induced plastic responses in Drosophila simulansfollowing exposure to combinations of temperature and humidity levels. J. Exp. Biol. 216, 4601-4607.

Cavicchi, S., Guerra, D., La Torre, V. and Huey, R.B.(1995). Chromosomal analysis of heat-shock tolerance in D. melanogaster evolving at different temperatures in the laboratory. Evolution 49, 676-684.

Chen, Z., Cuin, A.T., Zhou, M., Twomey, A., Naidu, P.B.Shabala, S(2007). Compatible solute accumulation and stress mitigating effects in barley genotypes contrasting in their salt tolerance. J. Exp. Bot. 58, 4245-4255.

Choudhary, N.L., Sairam, R.K. and Tyagi, A (2005). Expression of delta-l-pyrroline-5-carboxylate synthetase gene during drought in rice (Oryza sativaL.). Ind. J. biochem. Biophys. 42, 366-370.

Chown, S.L., Sorensen, J.G. and Terblanche, J.S. (2011). Water loss in insects: an environmental change perspective. J. Insect Physiol. 57, 1070-1084.

Colinet, H., Larvor, V., Laparie, M. and Renault, D. (2012). Exploring the plastic response to cold acclimation through metabolomics. Funct. Ecol. 26, 711-722.

Csonka, L.N. and Hanson, A.D. (1991). Prokaryotic osmoregulation:genetics and physiology. Annu. Rev. Microbiol. 45, 569-606.

Denlinger, D.L. and LeeR.E.Jr.(2010). Insect Low Temperature Biology.Cambridge Univ. Press, New York.

Fields, P.G., Fleurat-Lessard, F., Lavenseau, L., Febvay, G., Peypelut, L. and Bonnot, G.(1998). The effect of cold acclimation and deacclimation on cold tolerance, trehalose and free amino acid levels in Sitophilus granariesand Crypyolestes ferrugineus (Coleoptera). J. Insect Physiol. 44, 955-965.

Hoffmann, A.A. (2010). Physiological climatic limits in Drosophila: patterns and implications. J. Exp. Biol. 213, 870-880.

Hoffmann, A.A., Shirriffs, J. and Scott, M. (2005). Relative importance of plastic vs genetic factors in adaptive differentiation: geographical variation for stress resistance in Drosophila melanogasterfrom eastern Australia. Funct. Ecol. 19, 222-227.

Issartel, J., Renault, D., Voituron, Y., Bouchereau, A., Vernon, P. and Hervant, F. (2005). Metabolic responses to cold in subterranean crustaceans. J. Exp. Biol. 208, 2923-2929.

Kalra, B., Tamang, A.M. and ParkashR.(2017). Cross-tolerance effects due to heat hardening,desiccation and starvation acclimation of drosophilid-Zaprionus indianus. Comp. Bichem. Physiol. A. 209, 65-73.

Kaushik, J.K. and Bhatt, R (2003). Why is trehalose an exceptional protein stabilizer? J Biol Chem. 278, 26458-26465.

Kellermann, V., Heerwaarden, V.B., Sgro, C.M. and Hoffmann, A.A. (2009). Fundamental evolutionary limits in ecological traits drive Drosophilaspecies distributions. Science 325, 1244-1246.

Kellett, M., Hoffmann, A.A. and McKechnie, S.W. (2005). Hardening capacity in the Drosophila melanogasterspecies group is constrained by basal thermotolerance. Funct. Ecol. 19, 853-858.

Kemble, A.R. and MacPherson, H.T. (1954). Liberation of amino acids in perennial rye grass during wilting. Biochem. J. 58, 46-59.

Kostal, V., Zahradnickova, H. and Simek, P. (2011a). Hyperprolinemic larvae of the drosophilid fly,Chymomyza costata, survive cryopreservation in liquid nitrogen. Proc. Natl. Acad. Sci. U S A. 108, 13041-13046.

Kostal, V., Korbelova, J., Rozsypa, J., Zahradnickova, H., Cimlova, J., Tomcala, A., Simek, P.(2011b). Long-term cold acclimation extends survival time at 0 °C and modifies the Metabolomic profiles of the larvae of the fruit fly Drosophila melanogaster. PLoS One 6, 1-10, e25025.

Lee, R.E.(2010). A primer on insect cold tolerance. In: DenlingerD.L, LeeR.E., editors. Low temperature biology of insects. Cambridge: Cambridge University Press. pp. 3-34.

Liang, X., ZhangLu., Natarajan, S.K. and Becker, D.F. (2013). Proline mechanisms of stress survival. AntioxidRedox Signal. 19, 998-1011.

Malmendal, A., Overgaard, J., Bundy, J.G., Sorenson, J.G., Nielsen, N.C., Loeschcke, V. and Holmstrup, M. (2006). Metabolomic profiling of heat stress; hardening and recovery of homeostasis in Drosophila. Am. J. Physiol. 291, R-205-R-212.

Michaud, M.R. and Denlinger, D.L. (2007). Shifts in carbohydrates, polyols and amino acid pools during rapid cold hardening and diapause-associated cold hardening in flesh flies (Sarcophaga crassipalpis); a metabolomic comparison. J. Comp. Physiol. B. 177, 753-763.

Michaud, M.R., Benoit, J.B., Lopez-Martinez, G., Elnitsky, M.A., LeeR.EJr. and Denlinger, D.L. (2008). Metabolomics reveals unique and shared changes in response to heat shock, freezing, and desiccation in the Antarctic midge, Belgica antarctica. J. Insect Physiol. 54, 645-655.

Misener, S.R., Chen, C.P. and Walker, V.K. (2001). Cold tolerance and proline metabolic gene expression in Drosophila melanogaster. J. Insect Physiol. 47, 393-400.

Overgaard, J., Malmendal, A., Sørensen, J.G., Bundy, J.G., Loeschcke, V., Nielsen, N.C. and Holmstrup, M (2007). Metabolomic profiling of rapid cold hardening and cold shock in Drosophila melanogaster. J. Insect Physiol. 53, 1218-1232.

Parkash, R. and Munjal, A.K. (1999). Climatic selection of starvation and desiccation resistance in populations of some tropical Drosophilids. J. Zool. Syst. Evol. Res. 37, 195-202.

Parkash, R., Singh, S. and Ramniwas, S. (2009). Seasonal changes in humidity level in the tropics impact body color polymorphism and desiccation resistance in Drosophila jambulina- Evidence for melanism-desiccation hypothesis. J. Insect Physiol. 55, 358-368.

Parkash, R. and Ranga, P. (2014). Seasonal changes in humidity impact drought resistance in tropical Drosophila leontia: testing developmental affects of thermal versus humidity changes. Comp. Biochem. Physiol. A. 169, 33-43.

PiresC.S.S., Sujii, E.R., FrontesE.M.G., Tauber, C.A. and Tauber, M.J. (2000). Dry-season embryonic dormancy in Deois flavopicta(Homoptera: Cecropidae): roles of temperature and moisture in nature. Envt. Entomol. 29, 714-720.

Reina-Bueno, M., Argandoña, M., Nieto, J.J., Hidalgo-García, A., Iglesias-Guerra, F., Delgado, M.J. and Vargas, C (2012). Role of trehalose in heat and desiccation tolerance in the soil bacterium Rhizobium etli. BMC Microbiol. 12, 207-223.

Renault, D., Bouchereau, A., Delettre, Y.R., Hervant, F. and Vernon, P. (2006). Changes in free amino acids in Alphitobius diaperinus (Coleoptera: Tenebrionidae) during thermal and food stress. Comp. Biochem. Physiol. A. 143, 279-285.

Schmidt-Nielsen, K.(1990). Animal Physiology: Adaptation and Environment. 4th edn. Cambridge University Press,Cambridge.

Seymour, J.E. and Jones, R.E. (2000). Humidity-mediated diapauses in the tropical braconid parasitoid *Microplitis demolitor*. Ecol. Entomol. 25, 481-485.

Sgro, C.M., Overgaard, J., Kristensen, T.N., Mitchell, K.A., Cockerell, F.E. and HoffmannA.A,(2010). A comprehensive assessment of geographic variation in heat tolerance and hardening capacity in populations of Drosophila melanogaster from eastern Australia. J. Evol. Biol. 23, 2484-2493.

Shimada, K. and Riihima, A. (1990). Cold-induced freezing tolerance in diapausing and non-diapausing larvae of Chymomyza costata (Diptera: Drosophilidae) with accumulation of trehalose and proline. Cryo. Letters 11, 243-250.

Sinclair, B.J. (2015). Linking energetics and overwintering in temperate insects. J. Therm. Biol. 54, 5-11.

Sinclair, B.J., Ferguson, L.V., Salehipour-shirazi, G. and MacMillan, H.A. (2013). Cross tolerance and cross-talk in the cold: relating low temperatures to desiccation and immune stress in insects. Integr. Comp. Biol. 53, 545-556.

Szabados, L. and Savoure, A. (2009). Proline: a multifunctional amino acid Trends in Plant Sci. 15, 89-97.

Tauber, M.J., Tauber, C.A. and Masaki, S (1986). Seasonal adaptations of insects. Oxford University press.

Tauber, M.J., Tauber, C.A., Nyrop, J.P. and Villani, M.G. (1998). Moisture, a vital but neglected factor in the seasonal ecology of insects: hypotheses and tests of mechanisms. Environ. Entomol. 27, 523-530.

Tamang, A.M., Kalra, B. and Parkash, R. (2017). Cold and desiccation stress induced changes in the accumulation and utilization of proline and trehalose in seasonal populations of Drosophila immigrans. Comp. Biochem. Physiol. A. 203, 304-313

Verbruggen, N. and Hermans, C. (2008) Proline accumulation in plants: a review. Amino Acids 35, 753-759

Whitman, D.W. and Ananthakrishan, T.N. (2009). Phenotypic plasticity of insects: mechanisms and consequences. Science Publishers, Enfield, USA.

Williams, B.R., Heerwaarden, V.B., Dowling, D.K. and Sagro, C.M. (2012). A multivariate test of evolutionary constraints for thermal tolerance in Drosophila melanogater. J. Evol. Bio. 25, 1415-1426.

Williams, C.M., Watanabe, M., Guarracino, M.R., Ferraro, M.B., Edison, A.S., Morgan, T.J., BoroujerdiA.F.B. and Hahn, D (2014). Cold adaptation shapes the robustness of metabolic networks in Drosophila melanogaster. Evolution 68, 3505-3523.

Yancey, P.H., Clark, M.E., Hand, S.C., Bowlus, R.D. and Somero, G.N. (1982). Living with water stress: evolution of osmolyte systems. Science 217, 1214-1222.

